# Qualitative and quantitative bioanalytical methods validation of aristolochic acid DNA adducts and application in human formalin-fixed paraffin-embedded hepatocellular carcinoma tissues

**DOI:** 10.1101/2020.08.20.260331

**Authors:** Wang xin, Song jinping, Dong yapping, Hu linghua, Chen xiaoying, Yang fangfang, Qi haiyang, Qi xinming, Wen wen, Chen shuzhen, Xing guozhen, Ren jin

## Abstract

Mutation signature of aristolochic acid (AA) found in urothelial or hepatocellular carcinoma causes public concern about the cancer risk of AA. In contrast, direct evidence based on the reliable bioanalytical method for the exposure of AA is still lacking and not universal. Here, we strictly complied with the qualitative and quantitative guidance for forensic toxicological analysis: In the sample preprocessing, DNA from formalin-fixed and paraffin-embedded (FFPE) tissues was digested to single nucleotide by a series of enzymes with 70% enzymatic digestion efficiency. After protein precipitation, the samples were submitted to an ABI6500+ mass spectrometer for LC-MS/MS analysis. Ion pairs 543.2/427.2 and 543.2/395.2 of dA-AAI were selected from 5 ion pairs due to their higher LC-MS/MS response. Both these ion pairs have excellent selectivity and specificity in rat liver DNA matrix, and a linear regression range from 5 pg/mL to 200 pg/mL with the best fit and determination coefficient (r) greater than 0.99. The intra and inter batch accuracy and precision of these two ion pairs are also acceptable with less than 15% variation. The total recovery for ion pair 543.2/427.2 and 543.2/395.2 of dA-AAI was 90.06% and 90.76%, respectively. Our method has a minor matrix effect and good stability under different temperature and time conditions. With signal to noise ratio ≥ 3, 2 ion pairs (< 50 % relative abundance variation), the lower limit of quantification (LLOQ) of our method is set to 5 pg/mL(∼3.6 AAI-DNA adducts per 10^8^ DNA bases). By using this validated bioanalytical method of dA-AAI, 165 human HCC FFPE tissues were analyzed, the total ratio of samples with peak^543.2/427.2^ is 17.0% (28/165), with peak^543.2/395.2^ is 9.09% (15/165) which yields the total ratio of samples combined peak^543.2/427.2^ and peak^543.2/395.2^ is 7.27% (12/165). Two samples are higher than 5 pg/mL under the qualitative requirements. In conclusion, we first reported a fully validated methods to analyze the DNA adducts level of aristolochic acid, which could be qualitatively and quantitatively applied to the investigation of AA exposure in the human and other species.

## 1. Introduction

A recent whole-exome sequencing study linked aristolochic acids, a urothelial carcinogen, with the prevalence of hepatocellular carcinoma (HCC) in Asia, which showed 47% HCCs in China may be associated with the exposure of aristolochic acids^1-3^. Aristolochic acids (AA) contribute to the global prevalence of chronic kidney disease and urothelial cancer^4-6^. AA-formed mutagenic lesions, dA-aristolactam adducts (dA-AAI, Figure 1), produce a unique **AT-TA transversions** mutation at adenine mainly on the non-transcribed strand (**NTS strand bias**) with a notable peak at 5′-CTG-3′ **(trinucleotide contexts characteristic)**. These three characteristics constitute the mutational signature of AA^1-3, 5, 7^. Based on this **signature**, several studies have reported a correlation between AA and HCC with a coefficient from 25% to 47%^2, 8^. Every year, nearly 400, 000 Chinese people die from HCC, and 24 herb medicines from the *Aristolochiaceae* family are still being prescribed in the clinic^2, 9, 10^. However, direct exposure evidence of AA based reliable bioanalytical method is still lacking.

**Figure 1.**
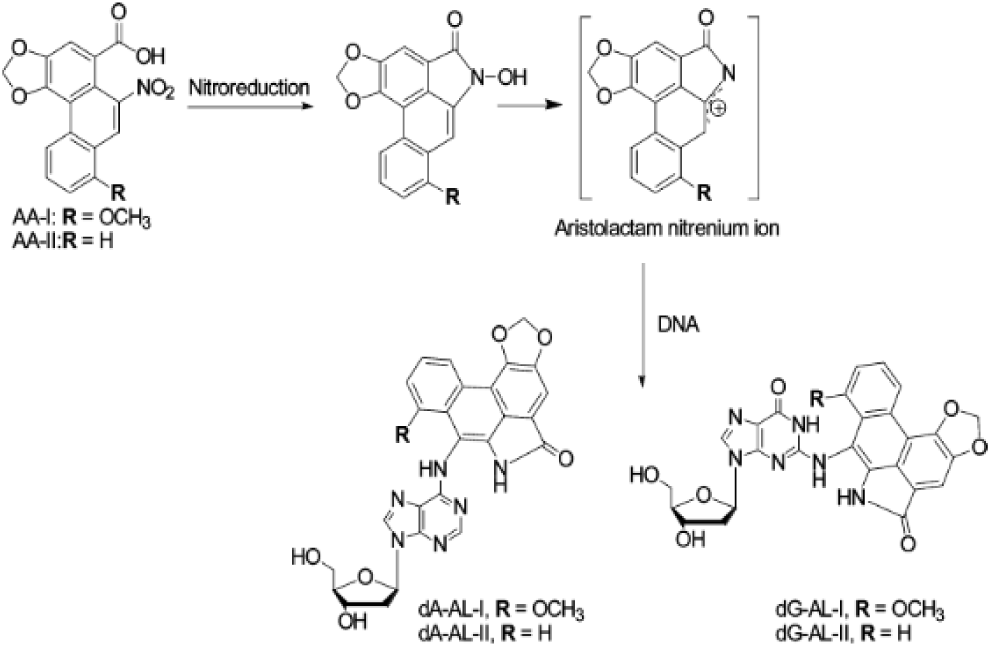
Metabolic activation and DNA adduct formation of aristolochic acid.

Herbs are widely used among Chinese people for disease treatment and diets. In the South of China, drinking herbal tea is a part of Cantonese lifestyle, over 1600 types of herbs are used in herbal tea, slow-cooker soup, or planted as homegrown herbs *Houttuynia cordata, Aristolochia cinnabarina* from *Aristolochiaceae* family^11-13^. The potential exposure of AA has aroused the tremendous public concerns in these provinces since 2017. It is emergent to investigate the risk of AA exposure in the population, and DNA adducts formed by AA are the best biomarker and direct evidence for AA exposure. At the same time, amounts of formalin-fixed and paraffin-embedded (FFPE) tissues archived in the hospitals provide an abundant sample resource for the population investigation of AA exposure.

The investigation of AA exposure will disclose the risk faced by the individual and may change the lifestyle of the population. Therefore, to avoid undue public alarm, we should be cautious to select bioanalytical methods and criteria to perform the qualitative confirming and quantitative test of DNA adducts in human samples, especially when the content is close to the lower limit of quantitation (LLOQ). Here, we strictly complied the guidance of doping control and forensic toxicological analysis, which strongly require reliable qualitative confirming for judicial expertise, and validated our bioanalytical methods with the following criteria and protocols: 2 ion pairs with < 50% relative abundance variation, signal to noise ratio ≥ 3 and quantitative validation protocols. With this validated method, we detected the potential of AA exposure in 165 HCC patients. Since the dA-AAI is the most dominant and persistent AA-DNA adduct ^1-3^, and it is the key AA-DNA adduct to generate A→T transversion mutation signature, hereby we only took this in count in this study.

## 2. Experimental

### 2.1 Chemicals and reagents

Aristolochic acid I (AAI, 99.63%) and aristolochic acid II (AAII, 98.8%) were purchased from Nanjing Spring & Autumn Biological Engineering Co., Ltd. 7-(deoxyadenosin-N6-yl) aristolactam I (dA-ALI) was synthesized based on Suzuki-Miyaura coupling reaction. The final coupling step that produced the adduct and ^1^H- NMR spectrum was reported in Supporting Information. Protease K, deoxyribonuclease (DNase I), alkaline phosphatase (AP), nuclease P1 (NP1), and phosphodiesterase I (PDI) were purchased from Sigma Aldrich (Shanghai, China). Zymo FFPE DNA miniprep kit was purchased from zymo Research (Beijing). ACS reagent grade formic acid (98%), isopropanol, β-mercaptoethanol, methanol, xylene, 75% ethanol, 95% ethanol, anhydrous ethanol, glacial acetic acid, hydrochloric acid, deoxyadenosine (dA), deoxyguanosine (dG), deoxycytosine (dC), deoxythymine (dT), and uracil (U) were purchased from SINOPHARM GROUP Co., Ltd. Methanol and acetonitrile were purchased from Merck.

### 2.2 Chromatographic conditions

A Sciex ExionLC™AD ultra-high-performance liquid chromatography (UHPLC) instrument was used in this study. The separation was carried out on an ACE C18 column (50 × 2.1 mm, 5 microns) maintained at 40°C. A gradient program was conducted using an aqueous mobile phase A of 0.2% acetic acid in water and an organic mobile phase B of acetonitrile. The flow rate was set at 0.5 mL/min through each injection. The gradient program was designed as shown in **Table 1**. The injection volume was 15 μL, and the retention time of the dA-AAI (Analyte) and AAII (IS) were 1.19 min and 1.42 min, respectively.

**Table 1.** The gradient elution procedure of this method

| Time<br>(min) | Flow rate<br>(mL/min) | Mobile phase A<br>(%) | Mobile phase B<br>(%) |
| --- | --- | --- | --- |
| 0.00 | 0.5 | 70 | 30 |
| 0.20 | 0.5 | 70 | 30 |
| 1.50 | 0.5 | 0 | 100 |
| 2.00 | 0.5 | 0 | 100 |
| 2.01 | 0.5 | 90 | 10 |
| 2.20 | 0.5 | 90 | 10 |
| 2.21 | 0.5 | 0 | 100 |
| 2.70 | 0.5 | 0 | 100 |
| 2.71 | 0.5 | 30 | 70 |
| 4.00 | 0.5 | 30 | 70 |

### 2.3 Mass spectrometry conditions

Detection was carried out by a Sciex TRIPLE QUAD® 6500 Plus MS/MS fitted with electrospray ionization (ESI) probe and operated in the positive ion mode. The detection was carried out in multiple reactions monitoring (MRM) mode. The optimized conditions were as follows: Curtain Gas, 40 psi; Collision Gas, 10 psi; IonSpray Voltage, 5000 V; Temperature, 550 V; Ion Source Gas 1, 50 psi; Ion Source Gas 2, 50 psi; Entrance Potential, 10 V; Collision Cell Exit Potential, 15 V. The MRM transitions and the related optimized declustering potential (DP), collision energy (CE) for dA-AAI and IS are shown in **Table 2**. Two ion pairs of the dA-AAI were chosen for the method validation.

**Table 2.** Mass spectrometry parameters

| Ion pairs | Q1<br>(Da) | Q3<br>(Da) | Time<br>(ms) | DP<br>(V) | CE<br>(V) |
| --- | --- | --- | --- | --- | --- |
| dA-AAI | 543.2 | 427.2 | 80 | 66 | 31 |
| dA-AAI | 543.2 | 395.2 | 80 | 55 | 65 |
| IS | 312.1 | 268.0 | 80 | 80 | 11 |

### 2.4 Preparation of standard and sample solutions

The standard stock solution of dA-AAI 1 mg/mL (w/v) was prepared with DMSO, while the standard stock solution of IS 1 mg/mL (w/v) was prepared with methanol. The dA-AAI working solutions for calibration and quality control samples were prepared from the stock solution by DMSO, and the IS working solution (1.00 μg/mL) was prepared by diluting the stock solution with ethanol.

### 2.5 Test sample preparation

#### 2.5.1 Tissue digestion stage

Two pieces of 10 μm sections with liver tissue were cut by the paraffin section machine, put into 1.5mL EP tube, and centrifuged at 10000 × g for 30s. 400 μL dewaxing agent was added into each 1.5ml EP tube, and then the samples were incubated at 55 °C for 1 min for dewaxing. 100 μL enzymolysis digestive juice (DEPC water 45 μL, 2 × digestion buffer 45 μL and protease K buffer 10 μL) was added into the dewaxed tissues, and then the samples were incubated in the water bath at 55 °C overnight for 16 hours for digestion. RNase A solution was added into the digested tissue samples and mixed well immediately after the samples were transferred and incubated in a metal bath at 94 °C for 20 min, and then let the samples stand at room temperature for 5 min.

#### 2.5.2 DNA purification stage

The 350 μL Genomic Lysis Buffer and 135 μL isopropanol was added into the digested tissue samples, and then centrifuged at 12000 × g for 1 min. The supernatant was transferred to a small column with a collecting tube (Zymo-Spin(tm) IIC Column), and centrifuged at 10000 × g for 1 min. 400 μL genomic DNA wash 1 was added into the small column after the collecting tube of the small column was replaced and centrifuged at 10000 × g for 1 min. 700 μL genomic DNA wash 2 was added into the small column after the waste liquid in collecting tube was removed and centrifuged at 10000 × g for 1 min. 200 μL genomic DNA wash 2 was added into the small column after the waste liquid in collecting tube was removed and centrifuged at 10000 × g for 1 min. 50μL DNA extraction buffer was added into the small column after the small column was transferred to a clean 1.5ml EP tube and centrifuged at 17000 × g for 5 min after stand for 5 minutes. The purified DNA in EP tube was eluted.

#### 2.5.3 Incubation stage

The 48 μL purified DNA was used for the incubation stage. 1.5 μL DNase I (Type IV from bovine pancreas; 2542 U/mL in 0.15M NaCl; 254.2 U/mg DNA) was added into the purified DNA, and the mixture was incubated at 37 °C for 1.5h. Next, 1 μL nuclease P1 (from Penicillium citrinum; 100U/mL in 1 mM ZnCl_2_; 4 U/mg DNA) was added into the mixture, and the incubation was continued at 37 °C for a further 3h. The 1.5 μL Alkaline phosphatase (from E. coli; 24 U/mL in 1mM MgCl_2_; 2 U/mg DNA) and 8 μL phosphodiesterase I (from Crotalus adamanteus venom; 1.7 U/mL in 110 mM Tris-HCl at pH 8.9 containing 110 mM NaCl, 15 mM MgCl_2_, and 50% glycerol; 0.0714 U/mg DNA) were added last, and the incubation was continued at 37 °C for additional 18 h.

#### 2.5.4 Protein precipitation stage

60 μL IS working solution was added into 60 μL incubation sample for protein precipitation and centrifuged at 12000 rpm for 5 min after full vortex. 110 μL supernatant was transferred to a 96-well plate for LC-MS/MS analysis.

### 2.6 Calibration and QC sample preparation

Six μL of dA-AAI working solutions were added to 54 μL drug-free SD Rat liver DNA matrix (prepared according to 2.5.1-2.5.3) to obtain dA-AAI concentration levels of 5, 10, 20, 50, 75, 100, 150 and 200 pg/mL respectively. Quality control (QC) samples were prepared at concentrations of 5 pg/mL (LLOQ), 15 ng/mL (LQC), 80 ng/mL (MQC) and 160 ng/mL (HQC). All the calibration and QC samples were then prepared as 2.5.4 for LC-MS/MS analysis.

### 2.7 Method validation

#### 2.7.1 Selectivity and Specificity

To verify the absence of interfering endogenous substances around the retention time of dA-AAI and IS, the selectivity and specificity of the method were investigated by chromatograms obtained from SD Rat liver DNA matrix, which contained no dA-AAI and IS.

#### 2.7.2 Linearity

dA-AAI calibration curves were obtained by plotting the peak area ratio (dA-AAI/IS) against concentrations of the calibrators. For the calibration curve, following concentrations of dA-AAI were used: 5 (3.6 adducts/10^8^ dN), 10 (7.2 adducts/10^8^ dN), 20 (14.4 adducts/10^8^ dN), 50 (36.0 adducts/10^8^ dN), 75 (54.0 adducts/10^8^ dN), 100 (71.9 adducts/10^8^ dN), 150 (107.9 adducts/10^8^ dN), 200 (143.9 adducts/10^8^ dN) pg/mL. The results were obtained using linear regression analysis, with the weighting factor being 1/x^2^.

#### 2.7.3 Accuracy and precision

Intra assay precision and accuracy of dA-AAI were calculated at LLOQ (5 pg/mL), LQC (15 pg/mL), MQC (80 pg/mL) and HQC (160 pg/mL) levels for six replicates, each of the same analytical run. Inter-assay precision and accuracy were calculated after the replicates in three different analytical runs.

#### 2.7.4 Recovery

The recovery (RE) of dA-AAI was calculated by comparing the peak area of the dA-AAI from the extracted sample with that obtained from an unextracted sample at the same concentration for the QC samples containing 15, 80, 160 pg/mL for dA-AAI. IS recovery was investigated by comparing all of the extracted and unextracted samples.

#### 2.7.5 Matrix effect

The matrix effect was evaluated according to the precision of the IS normalized MF of LQC and HQC samples. Extracted and aqueous samples were compared to determine the matrix factor (MF) for dA-AAI and IS. IS normalized MF was calculated by comparing dA-AAI MF with IS MF.

#### 2.7.6 Stability

Stability experiments were performed in triplicates of LQC and HQC samples. Freeze-thaw stability was evaluated after subjecting the QC samples to the freeze-thaw cycle from −80°C to room temperature 3 times. Benchtop stability was evaluated after subjecting the QC samples at room temperature for 6h. Autosampler stability was evaluated after the QC samples were left in autosampler set at 8 °C for 21h. Cryopreservation stability was evaluated after subjecting the QC samples at −80 °C condition for 7d. Samples were considered stable if the average measured concentration was within ±15% compared with the theoretical concentration.

#### 2.7.7 Carryover

Carryover was determined by injecting blank samples after injecting ULOQ samples. The peak area of dA-AAI must be no bigger than 20% of dA-AAI in accompanying LLOQ, while the peak area of IS must be no bigger than 5% of IS in accompanying LLOQ.

## 3. Results and application

### 3.1 Method Validation

#### 3.1.1 Selectivity and Specificity

The selectivity and specificity were assessed by analyzing extracted samples of analyte at ULOQ without IS, IS sample at working concentration without dA-AAI, and blank sample without dA-AAI and IS. The peak area of both ion pairs observed at the retention time of dA-AAI was less than 20% of the accompanying LLOQ area (5 pg/mL). It was found that IS is not interfering with dA-AAI and vice versa. Representative chromatograms are shown in **Fig. 2**.

**Figure 2.**
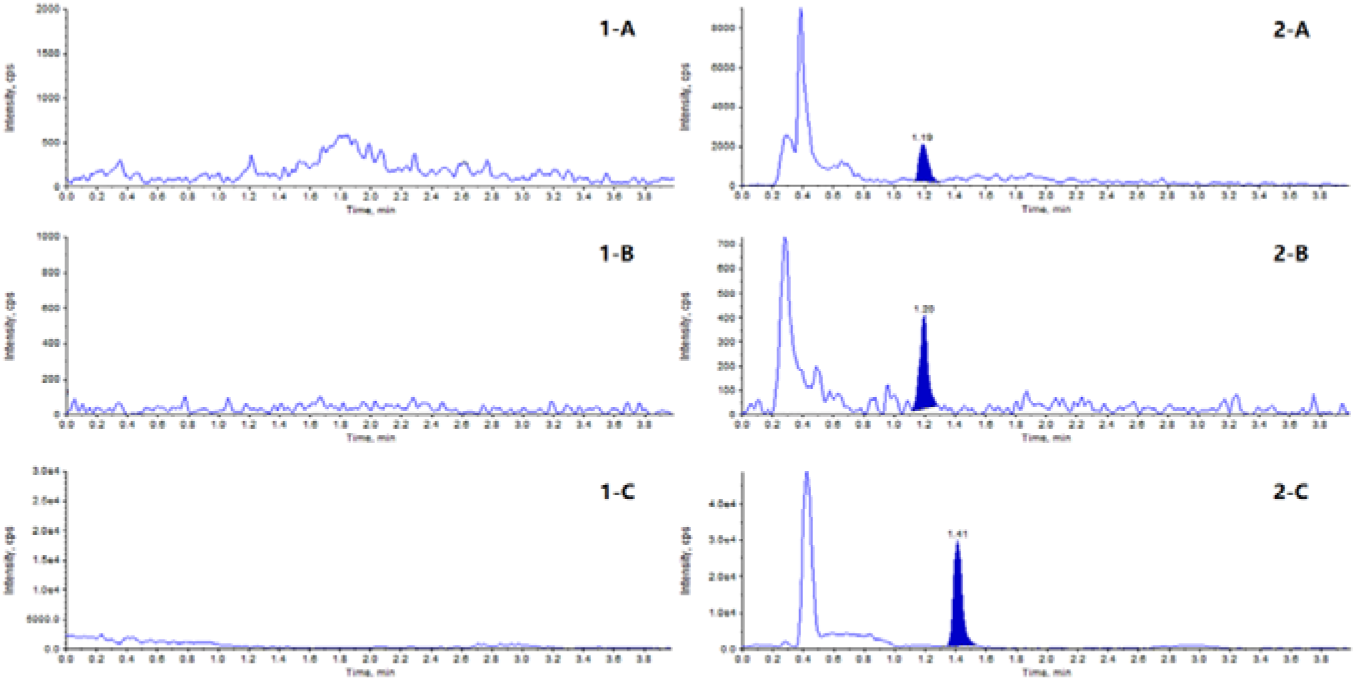
MRM chromatograms. Representative MRM chromatograms of dA-AAI(543.2/427.2) in (1-A) blank matrix; (2-A) matrix spiked with dA-AAI at LLOQ; Representative MRM chromatograms of dA-AAI (543.2/395.2) in (1-B) blank matrix; (2-B) matrix spiked with dA-AAI at LLOQ; Representative MRM chromatograms of IS in (1-C) blank matrix; (2-C) matrix spiked with IS.

#### 3.1.2 Calibration curve regression

The calibration curve regression for dA-AAI of both ion pairs was a linear regression range from 5 pg/mL to 200 pg/mL (weighting factor 1/x^2^). This gave the best fit and coefficient of determination (r) for validation and was higher than 0.99, which was in the acceptable range.

#### 3.1.3 Accuracy and precision

The Intra batch coefficients of variation ranged from 2.79 to 14.89%, and percentage accuracy ranged from 92.10 to 114.69% for ion pair 543.2/427.2 of dA-AAI. The Inter batch coefficients of variation ranged from 5.95 to 9.49%, and percentage accuracy ranged from 96.78 to 110.63% for ion pair 543.2/427.2 of dA-AAI. The Intra batch and Inter batch results of ion pair 543.2/427.2 are presented in **Table 2**.

The Intra batch coefficients of variation ranged from 4.23 to 18.18%, and percentage accuracy ranged from 92.23 to 115.87% for ion pair 543.2/395.2 of dA-AAI. The Inter batch coefficients of variation ranged from 6.47 to 15.31%, and percentage accuracy ranged from 101.80 to 107.43% for ion pair 543.2/395.2 of dA-AAI. The Intra batch and Inter batch results of ion pair 543.2/395.2 are presented in **Table 3**.

**Table 2.** dA-AAI intra and inter batch accuracy and precision of ion pair 543.2/427.2

| Theoretical Conc.<br>(pg/mL) |  | LLOQ | LQC | MQC | HQC |
| --- | --- | --- | --- | --- | --- |
|  |  | 5 | 15 | 80 | 160 |
| Inter A&P 01 | Accuracy (%) | 104.83 | 104.00 | 103.27 | 114.69 |
|  | CV (%) | 9.97 | 5.42 | 14.89 | 14.26 |
| Inter A&P 02 | Accuracy (%) | 92.10 | 100.44 | 102.75 | 108.13 |
|  | CV (%) | 7.13 | 7.02 | 5.76 | 2.51 |
| Inter A&P 03 | Accuracy (%) | 93.40 | 103.78 | 107.42 | 109.06 |
|  | CV (%) | 4.37 | 5.78 | 2.79 | 4.05 |
| Intra A&P | Accuracy (%) | 96.78 | 102.74 | 104.48 | 110.63 |
|  | CV (%) | 9.49 | 5.95 | 8.93 | 8.83 |

**Table 3.** dA-AAI intra and inter batch accuracy and precision of ion pair 543.2/395.2

| Theoretical Conc.<br>(pg/mL) |  | LLOQ | QC 3 | QC 2 | QC 1 |
| --- | --- | --- | --- | --- | --- |
|  |  | 5 | 15 | 80 | 160 |
| Inter A&P 01 | Accuracy (%) | 115.87 | 106.11 | 102.00 | 109.79 |
|  | CV (%) | 5.51 | 7.03 | 9.31 | 8.45 |
| Inter A&P 02 | Accuracy (%) | 97.90 | 101.78 | 98.75 | 106.67 |
|  | CV (%) | 18.18 | 6.68 | 7.10 | 3.88 |
| Inter A&P 03 | Accuracy (%) | 92.23 | 104.11 | 104.65 | 105.83 |
|  | CV (%) | 11.18 | 11.47 | 4.23 | 6.72 |
| Intra A&P | Accuracy (%) | 102.00 | 104.00 | 101.80 | 107.43 |
|  | CV (%) | 15.31 | 8.34 | 7.15 | 6.47 |

#### 3.1.4 Recovery

The recovery of dA-AAI and IS was evaluated by comparing the mean peak area of six extracted LQC, MQC, and HQC samples to those of six appropriately diluted aqueous solutions. For the ion pair 543.2/427.2 of dA-AAI, the mean recovery values at LQC, MQC, and HQC sample levels are 85.81%, 86.13%, and 98.22%, respectively. The total mean recovery is 90.06%. For the ion pair 543.2/395.2 of dA-AAI, the mean recovery values at LQC, MQC, and HQC sample levels are 90.48%, 87.11%, and 94.69%, respectively. The total mean recovery is 90.76%. For the IS, the mean recovery values at LQC, MQC, and HQC sample levels are 85.40%, 86.13%, and 103.29%, respectively. The total mean recovery is 91.61%. The result shows that the method has a good recovery of both dA-AAI and IS.

#### 3.1.5 Matrix effect

The matrix effect was evaluated according to the precision of the IS normalized MF of LQC and HQC samples. For the ion pair 543.2/427.2 of dA-AAI, the precision of the IS normalized MF at LQC and HQC is 9.63% and 3.96%. For the ion pair 543.2/395.2 of dA-AAI, the precision of the IS normalized MF at LQC and HQC is 12.65% and 7.30%. The result shows that the method has good matrix effect of dA-AAI.

#### 3.1.6 Stability

The stability of the dA-AAI and IS in SD Rat liver DNA matrix under different temperatures and time conditions were evaluated. As shown in **Table 4** and **Table 5**, the results show that dA-AAI is stable.

**Table 4.** Stability results of dA-AAI (ion pair 543.2/427.2)

| Stability experiments | Stability duration and temperature | Theoretical Conc. (pg/mL) | Measured Conc. (pg/mL) | % Bias | %CV |
| --- | --- | --- | --- | --- | --- |
| Autosampler Stability-LQC | 21h at 8 °C | 15 | 14.30 | 95.33 | 4.37 |
| Autosampler Stability-HQC | 21h at 8 °C | 160 | 167.67 | 104.79 | 4.40 |
| Freeze-thaw cycles-LQC | 3<sup>rd</sup> cycle | 15 | 14.70 | 98.00 | 7.67 |
| Freeze-thaw cycles-HQC | 3<sup>rd</sup> cycle | 160 | 166.33 | 103.96 | 3.87 |
| Benchtop Stability-LQC | 6h at 20 °C | 15 | 13.67 | 91.11 | 1.84 |
| Benchtop Stability-HQC | 6h at 20 °C | 160 | 163.00 | 101.88 | 1.23 |
| Cryopreservation Stability-LQC | 7d at -80 °C | 15 | 14.87 | 99.11 | 3.18 |
| Cryopreservation Stability-HQC | 7d at -80 °C | 160 | 166.67 | 104.17 | 4.54 |
| Incubation Stability-LQC | 22.5h at 37 °C | 15 | 16.00 | 106.67 | 1.05 |
| Incubation Stability-HQC | 22.5h at 37 °C | 160 | 162.17 | 101.35 | 4.89 |

**Table 5.**
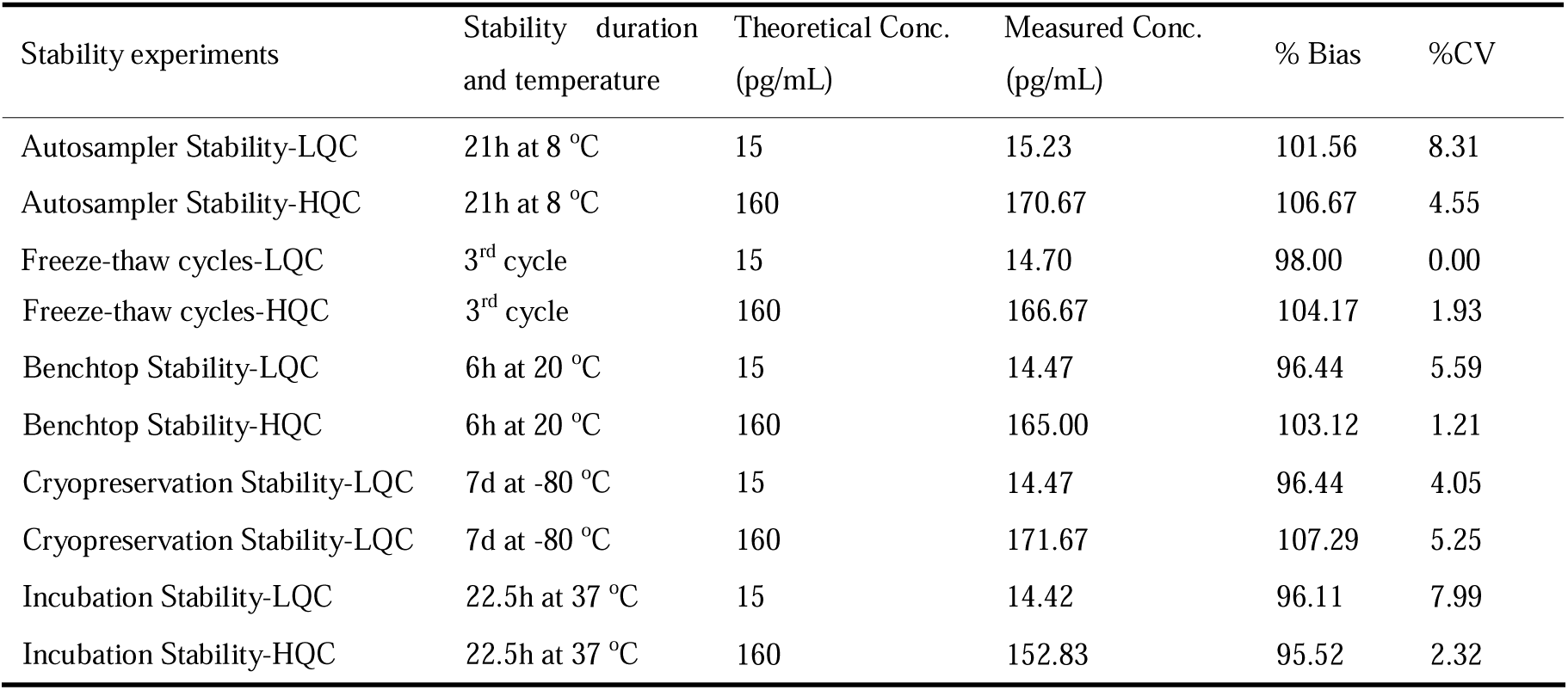
Stability results of dA-AAI (ion pair 543.2/395.2)

| Stability experiments | Stability duration and temperature | Theoretical Conc. (pg/mL) | Measured Conc. (pg/mL) | % Bias | %CV |
| --- | --- | --- | --- | --- | --- |
| Autosampler Stability-LQC | 21h at 8 °C | 15 | 15.23 | 101.56 | 8.31 |
| Autosampler Stability-HQC | 21h at 8 °C | 160 | 170.67 | 106.67 | 4.55 |
| Freeze-thaw cycles-LQC | 3<sup>rd</sup> cycle | 15 | 14.70 | 98.00 | 0.00 |
| Freeze-thaw cycles-HQC | 3<sup>rd</sup> cycle | 160 | 166.67 | 104.17 | 1.93 |
| Benchtop Stability-LQC | 6h at 20 °C | 15 | 14.47 | 96.44 | 5.59 |
| Benchtop Stability-HQC | 6h at 20 °C | 160 | 165.00 | 103.12 | 1.21 |
| Cryopreservation Stability-LQC | 7d at -80 °C | 15 | 14.47 | 96.44 | 4.05 |
| Cryopreservation Stability-LQC | 7d at -80 °C | 160 | 171.67 | 107.29 | 5.25 |
| Incubation Stability-LQC | 22.5h at 37 °C | 15 | 14.42 | 96.11 | 7.99 |
| Incubation Stability-HQC | 22.5h at 37 °C | 160 | 152.83 | 95.52 | 2.32 |

### 3.2 Qualitative confirming and quantitative analysis of dA-AAI in HCC samples

165 FFPE samples of HCC were obtained from Eastern Hepatobiliary Surgery Hospital with ethical approval and informed consent of the patients. Two 10 μm-thickness sections of each FFPE tissue were used for DNA isolation, adducts extraction, and dA-AAI analysis by LC-MS/MS.

Qualitative confirming was performed with a signal to noise ratio ≥ 3, two ion pairs with < 50% relative abundance variation (Table 6). In 165 samples, 12 samples have typical 543.2/427.2 and 543.2/395.2 peaks at 1.19 min, and 9 of 12 samples have calculated concentration of two ion pairs, and two samples have < 50% relative abundance variation between two ion pairs (Table 6, supplementary data 1).

**Table 6.** Qualitative confirming of dA-AAI in 165 HCC samples.

| Positive peaks of ion pairs | Signal to Noise ratio |  | % of HCC patients |  | Total ratio |
| --- | --- | --- | --- | --- | --- |
| | $\geq 3$ | $< 3$ | $\geq 3$ | $< 3$ | |
| 543.2/427.2 | 28 | 6 | 17.0% | 3.64% | 20.6% |
| 543.2/395.2 | 15 | 0 | 9.09% | / | 9.09% |
| 543.2/427.2 + 543.2/395.2 | 12 | 0 | 7.27% | / | 7.27% |
| < 50% Relative abundance variation | 2 | 0 | 1.21% | / | 1.21% |

Near the LLOQ, we performed quantitative validation at 1.23, 3.85, 5, 7.69 pg/mL, and finally set 5 pg/mL as the LLOQ due to the better stability, selectivity and accuracy/precision performance (Supplementary data 2, 3). With this method, we analyzed the dA-AAI level in 165 HCC samples. In 34 peak^543.2/427.2^ positive samples, 11 samples have calculated concentration ranged from 0-1 pg/mL, and two samples have calculated concentration bigger than 5 pg/mL. In 15 peak^543.2/395.2^ positive samples, 3 samples have calculated concentration ranged from 0-1 pg/mL, 9 samples have calculated concentration ranged from 1-5 pg/mL, and two samples have calculated concentration bigger than 5 pg/mL (The same one in peak^543.2/427.2^) (Table 7).

**Table 7.** Quantitative analysis of dA-AAI in 165 HCC samples.

| Ion pairs | Measured dA-AAI concentration (pg/mL) |  |  |  |
| --- | --- | --- | --- | --- |
| | $< 0$ | 0-1 | 1-5 | $> 5$ |
| 543.2/427.2 | 21 | 11 | 0 | 2 |
| 543.2/395.2 | 1 | 3 | 9 | 2 |

We also transformed the concentration of dA-AAI into the number of dA-AAI in 10^9^ nucleotides (dA-AAI/10^9^ dN). In all these 165 HCC samples, the numbers of dA-AAI/10^9^ dN are not more than 2 (Supplementary data 1).

## 4. Discussion

AA-DNA adducts detection is always the step since the AAs were found to be metabolically activated and bound to the DNA to generate the DNA adducts which yield the mutation and strongly related to the carcinogenesis. The ^32^P-postlabeling method was the most dominant way to determine the AA-DNA adducts since 1990s applied *in vitro* and *in vivo*^14^. However, because the radiation requirements, it was only limitedly used in the laboratory with radiation facility. The application of liquid chromatography-tandem mass spectrometry in determination of the AA-DNA adducts was a big step for AA-DNA adducts monitoring in which enables the ordinary lab could be involved. Chan et al reported that they used the liquid chromatography-electrospray ionization quadrupole time-of-flight mass spectrum (LC-ESI-qTOF MS) to determine the AA-DNA adducts in rat livers and kidneys with limitation is about 1/10^9^dN^14-15^. Grollman and his colleague found the LOQ value for the dA-AAI adduct is ∼0.3 adducts per 10^8^ DNA bases for UPLC-ESI/MS^3^, only using 10 μg DNA, and ∼0.3 adducts per 10^8^ for ^32^P postlabeling, using 20 μg DNA.

The UPLC-ESI/MS^3^ method is a superior method for the detection of AA-DNA adducts, particularly at trace levels ^16^. However, upon to date, there is no LC-MS AA-DNA adducts determination method was fully validated and no limitation value could be provided in human tissue especially when no enough evidence to demonstrate that human exposed to AA containing products. Hence, we modified the LC-MS method and proposed the limitation under our experiment condition which enable this method to have more application in lab and clinic.

We first reported a fully validated qualitative and quantitative methods to analyze the dA-AAI level in tissues. In our LC-MS/MS dA-AAI quantitative determination method, we explored the selectivity and specificity, linearity, accuracy and precision, recovery, matrix effect, stability, carryover, all are acceptable. In our method, we explored 5 μg DNA matrix in each sample, the limited determination level was 5 pg/mL. This level was equaled to the ∼3.6 adducts per 10^8^ DNA bases which was slightly higher than the results reported by Grollman and his colleague, but still in the same range^16^.

For the qualitative confirming, we referred to the guideline for quality control in forensic toxicological analyses from GTFCh: signal to noise ratio should be ≥ 3, and relative abundance variation between two ion pairs should be ≤ 50% (When one of product ion content is ≤10% that of another product ion in total ion current) ^17^. For quantitative measuring, we employed a standard quantitative validation protocol. Comparing to previous studies^14-16^, we designed dense concentration points near the LLOQ from 1.23 pg/mL to 7.69 pg/mL to fully validate the quantitative performance of our method, and finally, we set 5 pg/mL as our LLOQ (Supplementary data 2 and 3).

With this method, we qualitatively confirmed the potential exposure in 165 HCC samples. Two samples meet all criteria for qualitative judgment. In the quantitation measuring, the same two samples have a > 5 pg/mL dA-AAI level. However, we should carefully interpret this result, which could not reflect the actual situation of AA exposure. Firstly, each sample did not have the same DNA content (Supplementary data 1), and low DNA loading will reduce the positive ratio; Secondly, these 165 HCC samples were randomly selected from a tissue bank, and the clear follow-up information about herb using was absent. More tissues to isolate enough DNA, larger sample pools and definite history of herb using are necessary for the authentic exposure of AA in the population.

In conclusion, we first reported a fully validated methods to analyze the DNA adducts level of aristolochic acid, which could be qualitatively and quantitatively applied to the investigation of AA exposure in the human and other species.

## Acknowledgments

This work was supported by the National Science and Technology Major Project (2018ZX09101002-002).

